# Single-Molecule Imaging Reveals Differential Stability of Alpha-Synuclein Aggregates

**DOI:** 10.64898/2026.06.30.735576

**Authors:** Shih-Chin Wang, Stephanie J. Zhang, Tal Gilboa, John Kang, Anastasia Kuzkina, George T. Kannarkat, Liangxiao Chen, William M. Shih, Alice S. Chen-Plotkin, Vikram Khurana, David R. Walt

**Affiliations:** Department of Pathology, Brigham and Women’s Hospital, Boston, 02115, USA; Wyss Institute for Biologically Inspired Engineering, Harvard University, Boston, 02115, USA; Harvard Medical School, Boston, 02115, USA; Department of Neurology, Brigham and Women’s Hospital, Boston, 02115, USA; Department of Neurology, Perelman School of Medicine at the University of Pennsylvania, Philadelphia, 19104, USA; Department of Cancer Biology, Dana-Farber Cancer Institute, Harvard Medical School, Boston, 02115, USA; Department of Biological Chemistry and Molecular Pharmacology, Harvard Medical School, Boston, 02115, USA; Harvard Stem Cell Institute, Cambridge, MA 02138, USA; Broad Institute of Massachusetts Institute of Technology and Harvard, Cambridge, MA 02142 USA

## Abstract

α-Synuclein (α-syn) aggregation is central to Parkinson’s disease (PD), yet measurements in biofluids are confounded by the coexistence of monomeric and aggregated species. Using Syn-IMAGR, a single-molecule imaging platform with sub-femtomolar sensitivity, we show that purified α-syn aggregates undergo dilution-induced disassembly, revealing a concentration-dependent equilibrium. Applied to postmortem brain lysates, Syn-IMAGR distinguishes physiological α-syn multimers, which are dimmer and readily dissociate upon dilution, from PD-associated aggregates, which remain detectable and exhibit greater structural resistance to disruption. These results indicate that α-syn assemblies occupy distinct stability regimes, with PD-associated aggregates representing a more persistent and less dilution-sensitive structural state. Syn-IMAGR thus provides a quantitative framework for resolving α-syn species and for probing their concentration-dependent equilibrium.

## Introduction

Nearly one million people in the U.S. suffer from Parkinson’s disease (PD), a progressive neurological disorder that disrupts the nervous system and severely impairs daily life (*1, 2*). With approximately 90,000 people diagnosed with PD annually in the U.S. (*3*), PD is the second most common neurodegenerative disease (*4*). Despite available treatments that may slow disease progression, there remains an unmet need to diagnose PD early for timely intervention. To address this need, various diagnostic tools have been developed to detect PD and monitor disease progression (*5-10*). Among these detection platforms, biomarker-based molecular detection has gained significant traction due to its potential for non-invasive and high-sensitivity measurement. One of the most widely used biomarkers for PD diagnosis is α-synuclein (α-syn) (*11-14*). Aggregated α-syn has been studied in human cells, tissues, and biofluids for decades to investigate its role in PD pathogenesis and assess its potential as a reliable biomarker across various detection platforms (*15-18*).

Most antibody-based detection methods for α-syn quantification, such as enzyme-linked immunosorbent assays (ELISA) (*19-22*) face two main challenges. First, anti-α-syn antibodies fail to distinguish α-syn monomers, tetramers, and small aggregates (>200 kDa), leading to misinterpretation of aggregate composition. For example, in an assay targeting α-syn, a sample containing 50 α-syn monomers, 12 tetramers, or a single large aggregate composed of 50 monomers will yield vastly different signal intensities, even though each sample contains the same total number of α-syn molecules. Although antibodies have been developed to preferentially detect aggregated α-syn, these antibodies often retain some residual monomer binding at lower affinities (*23*). Given that monomeric, dimeric, and tetrameric α-syn species predominate in human biofluids (*24*), aggregation-specific antibodies can be saturated by non-pathological monomers, skewing quantitative assessments of pathological aggregates. Consequently, numerous studies have reported discordant α-syn measurements across platforms, highlighting a fundamental limitation of antibody-based detection.

Second, α-syn displays a unique feature: strong intermolecular interactions, rendering any aggregates resistant to dissociation. Unlike other proteins, α-syn aggregates remain stable even under high-stringency conditions, including treatment with detergents such as SDS and Triton X-100 (*23, 25*). As a result, strategies that attempt to improve detection by disassembling aggregates may be ineffective, since the aggregates largely retain their native conformation and resist antibody access. Detecting α-syn aggregates therefore requires far greater sensitivity than conventional assays provide. Standard immunoassays cannot resolve these rare assemblies from abundant monomers, making sub-femtomolar, single-molecule sensitivity essential for aggregates associated with PD.

To address these limitations, we developed Syn-IMAGR (α-Synuclein Imaging with Augmented Capture for Single-Molecule Granularity and Resolution), a single-molecule imaging platform that quantifies the fluorescence brightness of individual α-syn particles as a proxy for their molecular stoichiometry. This enables direct resolution of heterogeneous α-syn assemblies at sub-femtomolar sensitivity—overcoming the nanomolar detection limits of conventional single-molecule imaging (*26-28*). We systematically evaluated a panel of α-syn binders—including monoclonal antibodies, aggregate-selective antibodies, aptamers, and nanobodies—to identify reagents capable of reliably capturing aggregated species without high monomer background (*29*). Unexpectedly, screening and calibration experiments revealed that higher-order α-syn aggregates can disassemble into smaller species upon dilution, indicating that *in vitro*-generated α-syn aggregates behave as a reversible, concentration-dependent system rather than as a population of inert fibrils. This finding challenges long-held assumptions and suggests that sample handling may alter the measured aggregate composition. We also investigated whether similar dissociation behavior occurs in human samples.

Applying Syn-IMAGR to postmortem brain tissue and cerebrospinal fluid (CSF), we detected α-syn aggregates in both PD and neurological control brain lysates. Analysis of aggregate brightness revealed that PD brain lysates contain a higher proportion of larger aggregated α-syn relative to neurological controls. Here, we refer to the smaller aggregates detected in control brain lysates as physiological α-syn aggregates, reflecting assemblies that arise naturally in non-diseased human tissue, and to the larger aggregates enriched in PD brain lysates as PD-associated aggregates. PD-associated aggregates exhibited greater structural stability and reduced sensitivity to concentration-driven disruption compared to physiological α-syn aggregates from control tissue. In contrast, differences in CSF were much less pronounced, with aggregate concentrations at least 100-fold lower than those observed in brain lysates. We reason that α-syn concentrations in CSF are orders of magnitude lower than in brain tissue, such that physiological multimers likely dissociate below detection thresholds, compressing the dynamic range and making disease-associated differences difficult to resolve. This dilution-driven loss of detectable aggregates—rather than a distinct biological shift in CSF—likely explains the subtle differences observed between PD and control CSF. Together, these observations reveal fundamental features of α-syn assembly behavior in human samples for the first time and demonstrate how Syn-IMAGR overcomes key limitations of conventional detection methods while uncovering biases introduced by dilution and equilibrium dynamics.

## Results

### Characterization of α-Syn Standards: Skewed Detection Toward Monomeric α-Syn

Accurately distinguishing between monomeric and aggregated α-syn is critical for unbiased quantification in diagnostic and mechanistic studies. However, since α-syn aggregates are composed of many monomer units (**Fig. 1A**), conventional antibody-based assays report signal based on antibody binding events rather than on the number of α-syn molecules present. As a result, an aggregate containing many α-syn monomers can generate a signal comparable to—or lower than—that of a much smaller number of monomers, leading to biased measurements.

**Figure 1.**
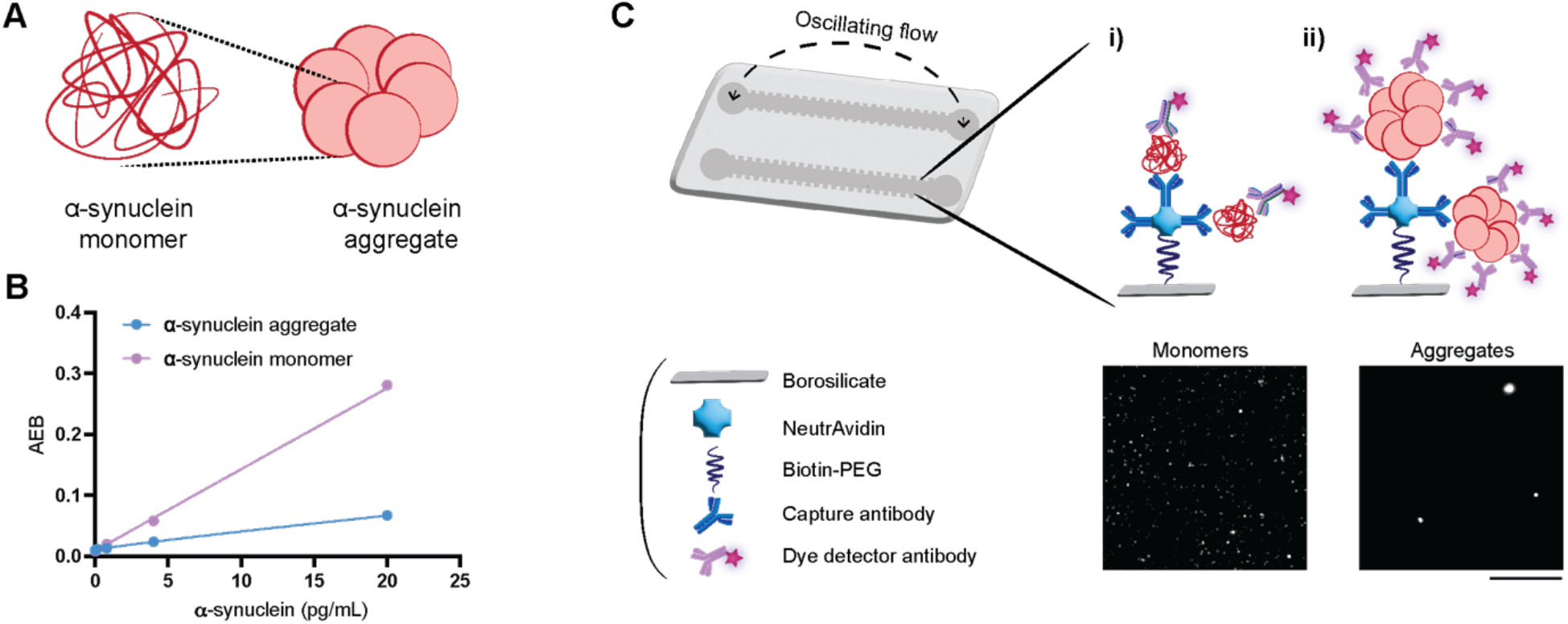
Single-molecule imaging of α-synuclein monomers and aggregates using Syn-IMAGR. (A) Schematic of α-synuclein monomers and their assembly into aggregates. (B) Calibration curves for monomeric (pink) and aggregated (blue) α-synuclein were generated using Single Molecule Array (Simoa) technology. Data are plotted as average enzyme per bead (AEB), representing the values of mean and standard derivation from duplicates. (C) The Syn-IMAGR platform employs a microfluidic device with oscillatory flow to enhance the capture of target proteins on a functionalized borosilicate surface coated with NeutrAvidin and immobilized biotinylated capture antibodies. Fluorophore-labeled detection antibodies bind to captured α-synuclein species. Monomers (i) generate low-intensity, compact fluorescent spots, while aggregates (ii) recruit multiple detection antibodies, yielding brighter and larger signals. Representative microscopy images highlight the distinct signal profiles. Scale bar, 10 µm.

To assess whether purified monomeric and aggregated α-syn standards (**Fig. S1**) used in immunoassays accurately reflect analyte composition, we quantified each species using an ultrasensitive single-molecule array (Simoa) assay. In this system, the average enzyme per bead (AEB) serves as a digital readout proportional to the number of antibody–antigen binding events. Notably, at the same total α-syn concentration, AEB values were consistently higher for monomeric samples than for aggregated samples comprising on the order of tens of monomers per assembly (e.g., ≥15 monomers, as estimated by Coomassie blue staininganalysis in **Fig. S1**) (**Fig. 1B**). The aggregated α-syn standard was characterized by Atomic Force Microscopy (AFM) to show the heterogeneity of α-syn species in the aggregate standard (**Fig. S2**). This disparity indicates that aggregated α-syn is undercounted relative to monomers, likely since the output signal amplified by an enzymatic reaction is counted above a particular threshold instead of signal levels in the antibody-based assay. As a result, an aggregate containing many α-syn monomers can generate a signal comparable to—or lower than—that of a much smaller number of monomers, leading to biased measurements. These results suggest that conventional antibody-based assays may systematically overestimate monomer abundance while underestimating aggregate content. This highlights a fundamental limitation of bulk immunoassays: the inability to resolve α-syn species with distinct sizes and structural conformations. These findings underscore the need for detection strategies capable of differentiating individual α-syn species to enable accurate compositional and quantitative analyses.

### Development of a Single-Molecule Imaging Assay for Species-Resolved α-Syn Detection

To resolve heterogeneous α-syn species in biofluids, we developed Syn-IMAGR, a single-molecule imaging platform with sub-femtomolar sensitivity and species-level resolution (**Fig. 1C**). This approach addresses the limitations of bulk immunoassays, which cannot distinguish between monomers, small aggregates, and higher-order aggregates commonly found in PD CSF. In this platform (**Fig. 1C**), we immobilized capture antibodies targeting α-syn on the surface of microfluidic channels, through which samples were continuously flowed to enhance binding efficiency by increasing target–surface interactions (*30*). Fluorescently labeled binders were then introduced to visualize α-syn molecules as discrete fluorescent spots, with spot intensity reflecting molecular size. To determine the optimal binding pair for distinctive α-syn species, we evaluated a panel of commercially available binding reagents under conditions enriched in either monomeric or aggregated α-syn (**Fig. 2**). Three monoclonal antibodies were tested as capture reagents: clones A and B, which recognize total α-syn, and clone C, which was developed to preferentially bind aggregated α-syn. Detection reagents included fluorophore-conjugated monoclonal antibodies, aptamers, and nanobodies (**Figs. 2 and S3**). The inclusion of aptamers and nanobodies enabled evaluation of whether smaller binders reduce steric hindrance and improve access to individual α-syn molecules within aggregated complexes.

**Figure 2.**
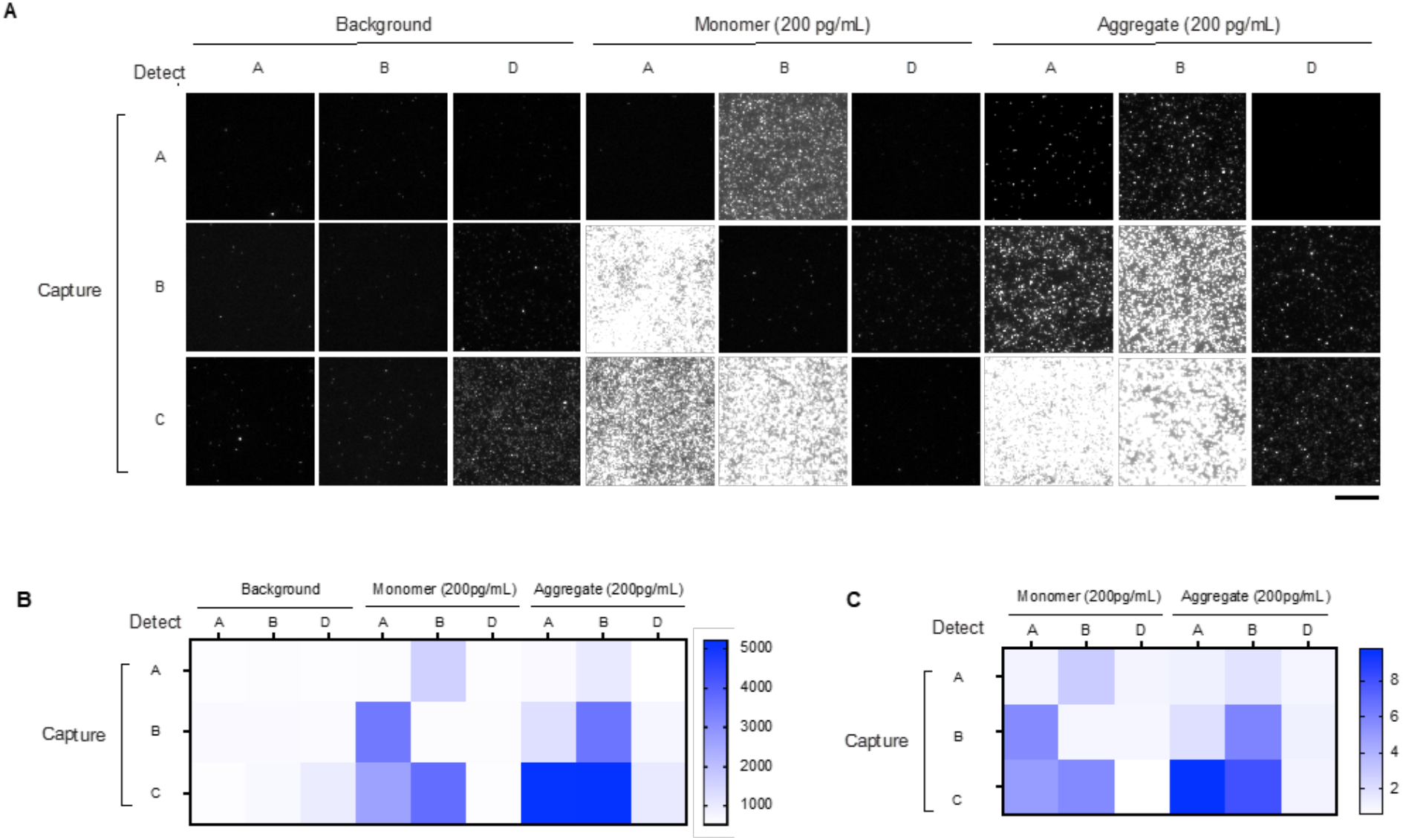
Validation of α-synuclein antibodies against distinct α-synuclein species using Syn-IMAGR. (A) Syn-IMAGR was used to assess antibody specificity against monomeric and aggregated α-synuclein (200 pg/mL) or buffer control. Capture was performed using α-synuclein antibodies (clones: A, B, and C), followed by detection with dye-conjugated α-synuclein antibodies (clones: A, B, and D). Representative images are shown with identical contrast settings. (B) The integrated fluorescence intensity of each condition was quantified and displayed as a heatmap. (C) Signal-to-noise ratios were calculated by dividing the integrated intensity of the monomeric or aggregated sample by that of the buffer control and presented as a heatmap. Scale bar: 10 µm.

Among the antibody pairs tested, A exhibits preferential binding to monomeric α-syn when paired with B as the detector. The B/A (capture/detector) configuration is more sensitive to monomer binding than the reverse pair. The B/B pair yielded low background in monomer-only conditions and higher signals in the presence of aggregates. Consistent with previous studies (*23, 31*), the aggregate-specific C still captured monomeric α-syn, but preferentially captured more aggregates, particularly when paired with A. In contrast, the binder F and nanobodies (D and E) failed to produce sufficient signal for sensitive detection, likely due to lower binding affinity or suboptimal epitope accessibility. Based on these results, we selected B/B and C/A pairs for subsequent calibration and quantitative analysis. Together, these two combinations offered complementary strengths: B/B provided high specificity for aggregated species with minimal background from monomers, while C/A offered higher specificity for aggregates and retained the ability to recognize other α-syn species.

### Dilution-Dependent Disassembly of α-Syn Aggregates Detected by B/B

Having validated the B/B antibody pair under monomeric conditions, where only minimal signal was detected, we proceeded to generate calibration curves for total and aggregated α-syn to assess whether this pair preferentially binds to aggregated species (**Fig. 3A**). Fluorescent spots were identified in each field of view (FOV), and the obtained mean number of spots per FOV was used as the readout for each measurement (**Fig. 3B**). When measuring total α-syn, the signal plateaued at approximately 100 spots per FOV, likely reflecting non-specific surface deposition of single detection antibody molecules (black dots in **Fig. 3B**). Of note, when purified aggregate standards were diluted to 0.16 pg/mL, we observed an increase in dimmer fluorescent spots (**Fig. 3A**, top), which corresponded to a distinct peak in the aggregate calibration curve (**Fig. 3B**). This observation suggests that larger aggregates disassemble to smaller species upon dilution at or above this concentration, altering the distribution of α-syn species in solution. These results imply that sample dilution could perturb the aggregate-monomer equilibrium, potentially introducing bias in other protein quantification assays. To selectively analyze larger aggregated species and remove background noise, we constructed a spot intensity histogram from blank samples, fitted it with a Gaussian distribution, and defined a threshold that eliminated 99.99% of background and smaller aggregates (**Fig. 3C**). Applying this threshold enabled us to isolate and quantify higher-order aggregates from both total and aggregated α-syn measurements. Notably, the B/B pair did not yield detectable signal in the total α-syn condition, consistent with prior reports that total α-syn preparations predominantly consist of monomeric, dimeric, and tetrameric species (*32*), which fall below the intensity threshold. The limit of detection for larger aggregates using the B/B pair was determined to be 0.29 pg/mL—comparable to the sensitivity achieved in other ultrasensitive protein assays (*21*).

**Figure 3.**
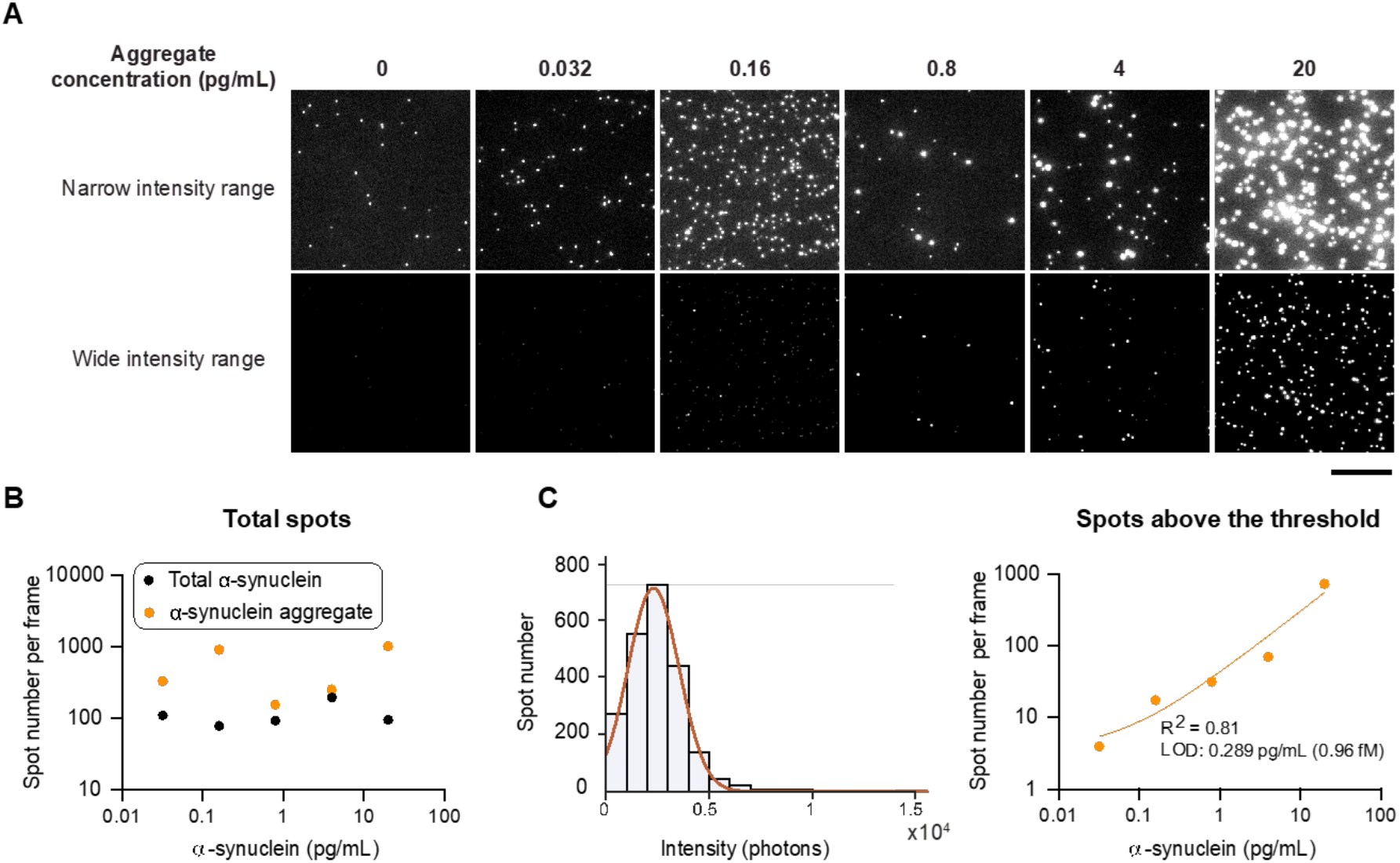
Calibration of α-synuclein aggregates using Syn-IMAGR with B/B antibody pair. (A) α-Synuclein aggregate and total α-synuclein standards were measured using Syn-IMAGR with the optimal capture/detection antibody pair (clone B). Representative images across the calibration range are shown with two different adjusted intensity ranges to highlight signal differences between monomeric and aggregated forms. (B) Bright spots per frame were counted and plotted against concentration for total α-synuclein (black dots) and aggregate (yellow dots). (C) Spot intensities from buffer-only controls were fitted with a Gaussian distribution (left panel) to define background from non-specific binding. After background subtraction, calibration curves were generated for total and aggregated α-synuclein. The calibration curve of total α-synuclein was not shown on a log y-axis since total α-synuclein was rarely captured by B/B antibody pair. Scale bar: 10 µm.

### C/A Antibody Pair Preferentially Detects Larger α-Syn Aggregates Due to the High Aggregate Affinity of C

In addition to the B/B pair, we evaluated the performance of the C/A antibody pair by generating calibration curves for both monomers and aggregates using Syn-IMAGR (**Fig. 4A**), building on the differential binding patterns of monomers and aggregates observed in **Fig. 2C**. For both monomeric and aggregated α-syn, total spot counts per FOV were used as the readout (**Fig. 4B**). Unlike the B/B pair, the C/A calibration curve for aggregates did not display a distinct peak at 0.16 pg/mL—suggesting that this pair is less sensitive to changes in small aggregate composition at lower concentrations (**Fig. 4C**). Consistent with previous studies (*23, 31*), the slope of the calibration curve for aggregates is notably shallower than that for monomers, demonstrating the preferential binding of the C/A pair to larger aggregated species. This suggests that once larger aggregates disassemble into smaller aggregates or monomers, the detection capability of the C/A pair may be compromised. Following background subtraction using Gaussian fitting of blank samples, the aggregate calibration curve was more sensitive than the monomer calibration curve (**Fig. 3C**), consistent with previous studies of enhanced recognition of higher-order aggregates by antibody C. In contrast, the B/B pair offers greater selectivity for aggregates in biofluids. While monomers may bind to the antibody B-coated surface, they are not effectively recognized by the antibody B detection antibody, thereby reducing background signal and enabling more accurate quantification of small aggregates and higher-order aggregates.

**Figure 4.**
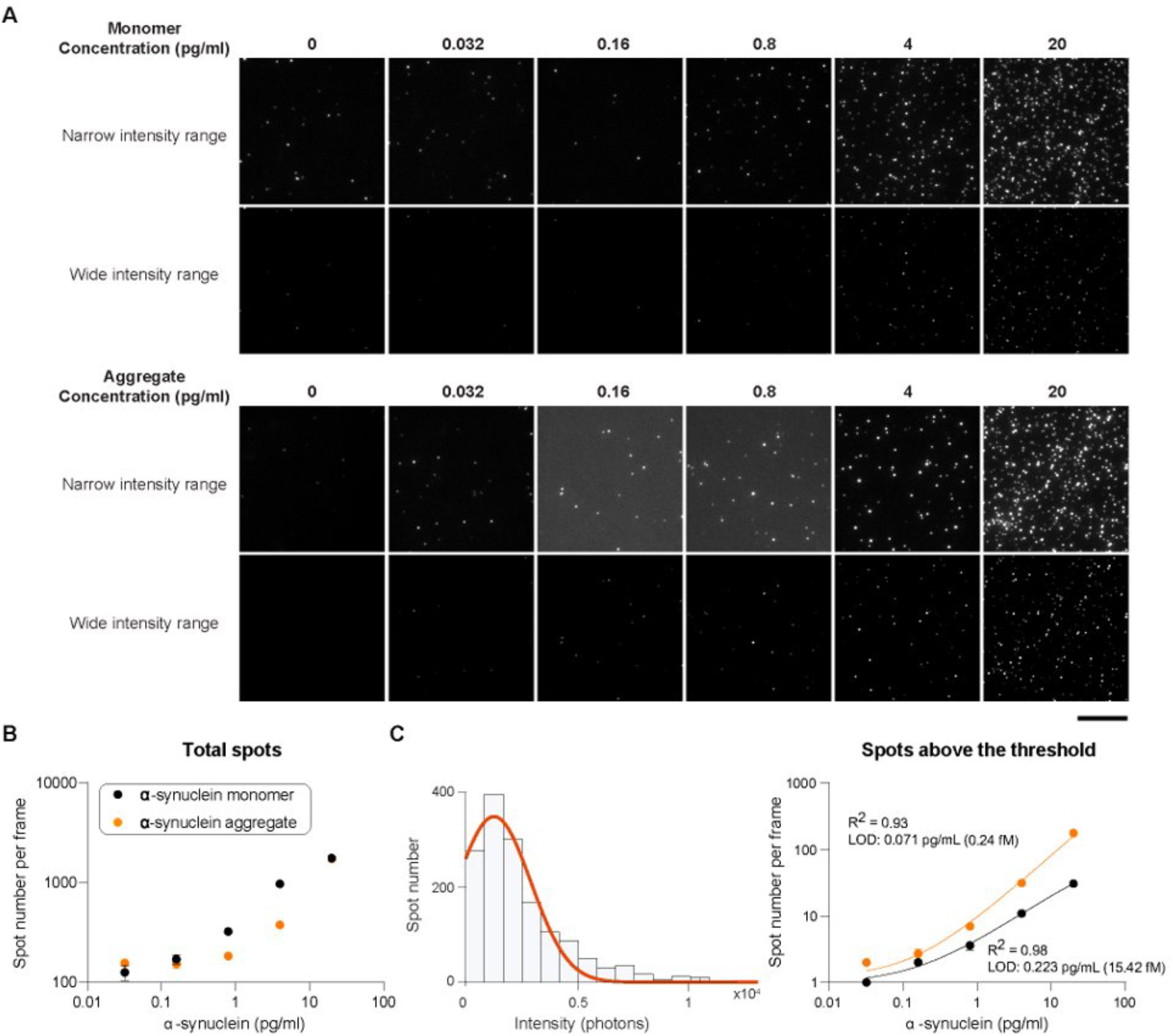
Calibration curve of α-synuclein monomers and aggregates using C/A antibody pair in Syn-IMAGR. (A) α-Synuclein monomer and aggregate standards were measured using Syn-IMAGR with clone C for capture and clone A for detection. Representative calibration images for monomer (top two rows) and aggregate (bottom two rows) across increasing concentrations. For each condition, images are displayed using two separate intensity ranges—a narrow range to enhance detection of dimmer spots and a wide range to highlight intensity differences among brighter signals. (B) Fluorescent spots were quantified per frame and plotted against concentration for monomer (black dots) and aggregate (yellow dots). (C) Spot intensities from buffer-only controls were fitted with a Gaussian distribution (left panel) to define non-specific background. After thresholding, calibration curves were generated for monomer and aggregate based on specific spot counts (black and yellow lines, respectively). Scale bar: 10 µm.

### Syn-IMAGR Reveals Distinct Equilibrium Behaviors of Physiological and PD-Associated α-Synuclein Aggregates

To minimize background from abundant monomeric α-syn in post-mortem brain lysates, we selected the B/B antibody pair, which preferentially recognizes aggregated and small assembled α-syn species. We reasoned that if PD pathology reflects a shift in α-syn homeostasis toward aggregation, then PD brain tissue should contain a higher proportion of aggregated species without necessarily altering total α-syn concentration. Using Syn-IMAGR, we visualized discrete α-syn fluorescent spots corresponding to individual aggregates and quantified both total and large assemblies using predefined intensity thresholds (**Fig. 5A**). PD brain lysates exhibited significantly higher aggregate counts than neurological controls, with large aggregates particularly enriched in PD (p = 0.0079; **Fig. 5B**). When normalized to total α-syn fluorescence, PD samples showed significantly elevated aggregate-to-total ratios (p = 0.0317 for total aggregates; p = 0.0079 for large aggregates; **Fig. 5C**), indicating that a larger fraction of the α-syn pool exists in aggregated form in PD tissue.

**Figure 5.**
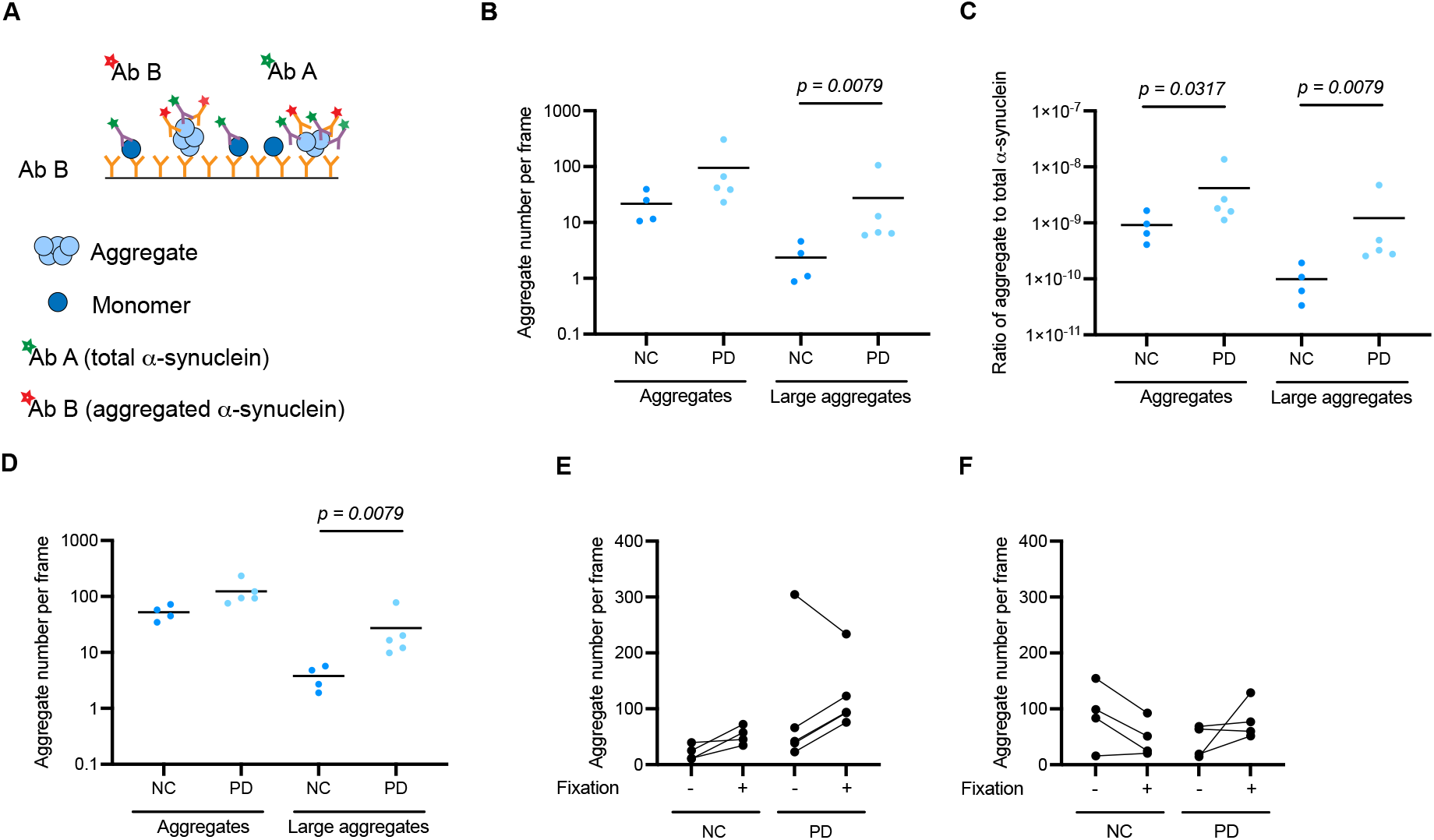
Measurement of α-synuclein species in brain lysates from neurological controls and PD patients in Syn-IMAGR. (A) The schematic diagram of two-color Syn-IMAGR detecting α-synuclein aggregates with antibody pair B/B and total α-synuclein with antibody pair B/A. (B) α-Synuclein aggregates in brain lysates from NC (n=4) and PD patients (n=5) at 100-fold dilution were measured using Syn-IMAGR with antibody pair B/B. The number of aggregates was counted based on different thresholds. Total aggregates (left) were determined with the same threshold as Figure 3C (mean±4xSD), whereas larger aggregates (right) were counted above higher threshold (mean ± 12 x SD). Large aggregates in PD brain lysates were increased significantly compared to brain lysates of NC. (C) Ratio of aggregates to total α-synuclein was calculated by dividing number of aggregates by integrated intensity of total α-synuclein. The ratio of either total aggregates or large aggregates over total α-synuclein was significantly higher in brain lysates from PD than NC. (D) α-Synuclein aggregates in brain lysates were fixed and diluted at 100-fold. Fixed aggregates were quantified based on the thresholds applied in (B). Increased levels of total aggregates and large aggregates were observed in brain lysates from PD. Total aggregates were compared in the absence and presence of fixation at 100-fold (E) and 1,000-fold (F) dilution. At 100-fold dilution, most brain lysates showed elevated numbers of total aggregates, whereas aggregate number in NC was reduced at 1,000-fold dilution. Mean of each group is shown as a dark line. The data were analyzed with a t-test.

We next asked whether these assemblies reflect a stable or reversible equilibrium by perturbing samples through serial dilution. In control samples, aggregate counts increased at 1,000-fold dilution (**Fig. S7A, D**), a counterintuitive rise that likely reflects the fragmentation of larger physiological multimers into multiple smaller assemblies that exceed the detection threshold. This behavior mirrors the disassembly pattern observed for synthetic aggregates at ∼0.16 pg/mL (**Fig. 3**), consistent with physiological multimers occupying a dilution-labile equilibrium. In contrast, PD samples showed decreased aggregate counts and reduced spot brightness at 1,000- and 10,000-fold dilutions (**Fig. S6–S7**), indicating that fewer PD-associated assemblies remain detectable at higher dilutions. Together, these data support a model in which physiological α-syn multimers (smaller, dilution-labile assemblies enriched in control brain lysates) exist as reversible, concentration-dependent species, whereas PD-associated aggregates (larger, more stable assemblies enriched in PD tissue) occupy a more stable and less dissociable region of the equilibrium landscape.

This interpretation aligns with reports that destabilization of the native α-helical tetramer favors formation of labile oligomeric intermediates in healthy tissue (*33*). Fixation experiments further reinforced this distinction. When α-syn assemblies were chemically fixed prior to dilution, aggregate counts increased in both groups, but the effect was substantially larger in controls (**Fig. 5D–E**). In control lysates, fixation preserved assemblies that would otherwise dissociate upon dilution, whereas in PD samples fixation produced only modest changes. At 1,000-fold dilution, cross-linking prevented disassembly of physiological multimers, reducing their apparent abundance relative to unfixed samples. To assess whether these equilibrium features extend systemically, we analyzed CSF from controls and PD patients. Aggregate counts were similar between groups, but PD CSF exhibited lower total α-syn and reduced aggregate-to-total ratios (**Fig. S7**). Given the high dilution of CSF relative to brain lysates, physiological multimers likely dissociate below detection thresholds, which may obscure subtle differences in aggregate abundance.

Collectively, these results with Syn-IMAGR resolve fundamentally distinct equilibrium behaviors of physiological and PD-associated α-syn assemblies (**Fig. S8**). Physiological multimers are dynamic, concentration-dependent, and dilution-labile, whereas PD-associated aggregates are more abundant, structurally stable, and resistant to dissociation, properties that may underlie their pathogenic persistence.

### Syn-IMAGR Detects Elevated Aggregated α-Syn in PD CSF

To minimize background signal from abundant monomeric α-syn in CSF, we chose the B/B pair for specific detection of α-syn aggregates and small aggregates. We hypothesized that patients with PD would exhibit elevated levels of α-syn aggregates, while monomer levels would remain comparable to those in neurological controls. However, dilution of CSF can perturb the equilibrium between monomers and aggregates, introducing bias in the measurement of aggregates or small aggregates. To address the challenge of detecting α-syn aggregates while simultaneously quantifying total α-syn levels, we employed a two-color Syn-IMAGR strategy. Aggregated α-syn was detected using the B/B antibody pair, while total α-syn was measured using dye-conjugated A, which broadly recognizes all α-syn species and reports integrated fluorescence intensity. We observed that total α-syn levels were significantly elevated in all three PD patients compared to controls, with PD1 and PD3 showing the most pronounced increases (*p* value < 0.0001) and PD2 showing a moderate but significant elevation (*p* value < 0.01) (**Fig. 5A**). PD patients 1 (2.7±0.4 pg/mL; *p* value < 0.0001) and 2 (24.5±1.6 pg/mL; *p* value < 0.0001) exhibited significantly elevated levels of aggregated α-syn compared to controls (**Fig. 5B**). Notably, PD 2 showed the highest level of aggregated α-syn among the samples we tested but displayed the lowest level of total α-syn among PD patients. This finding highlights the potential for conventional total α-syn measurements to obscure clinically relevant increases in aggregated species. Additionally, PD patient 3 (1.3±0.3 pg/mL; *p* value < 0.1, not significant) showed modestly elevated total α-syn levels relative to controls. Collectively, these results demonstrate that Syn-IMAGR enables precise, species-specific quantification of α-syn in human CSF, enabling more accurate assessment of disease-associated changes. This approach could potentially lead to improved PD diagnosis and progress monitoring through unbiased detection of aggregated α-syn.

## Discussion

We evaluated 15 capture/detection combinations for their ability to distinguish between α-syn monomers and aggregates. Based on sensitivity and specificity toward aggregated species, we selected the B/B pair to detect aggregates. To account for potential steric hindrance that may impair antibody access to epitopes on densely packed aggregates or small aggregates, we included two nanobodies, D and E, in our screening. Despite their smaller size, these nanobodies demonstrated suboptimal affinity for aggregated α-syn and exhibited elevated background signals in buffer-only conditions, precluding their effective use in sensitive single-molecule detection. Our findings also challenge the presumed aggregate-specificity of the C antibody. Although antibody C is often described as selective for aggregated α-syn, our results suggest that it binds both monomeric and aggregated forms, particularly smaller aggregates, which likely interferes with efficient detection by fluorophore-conjugated antibodies. This was evident in buffer-spiking experiments, where differential binding affinities to monomeric versus aggregated α-syn were readily observed. Together, these results establish Syn-IMAGR as a framework for resolving heterogeneous protein assemblies in complex biological samples with single-molecule sensitivity. Beyond α-synuclein, this approach may be broadly applicable to a wide range of aggregation-associated proteins implicated in neurodegenerative disease, including amyloid-β and tau in Alzheimer’s disease, TDP-43 and superoxide dismutase 1 in amyotrophic lateral sclerosis, mutant huntingtin in Huntington’s disease, and prion proteins in transmissible spongiform encephalopathies.

We next examined the physiological relevance of these observations using post-mortem brain lysates from PD patients and neurological controls. Syn-IMAGR directly visualized a pronounced enrichment of α-syn aggregates in PD tissue and a significantly elevated aggregate-to-total α-syn ratio. When these lysates were serially diluted, we observed two distinct behaviors: physiological aggregates fragmented and dissociated readily, while PD-associated aggregates maintained detectability and brightness far more consistently across dilution. Thus, α-syn assemblies are not static fibrils, but exist along a spectrum in which the degree of reversibility differs sharply between physiological and PD-associated species. These findings indicate that the aggregated state in PD represents a shift toward more stable, less dissociable assemblies, whereas physiological α-syn multimers occupy a highly dynamic, concentration-dependent equilibrium (**Fig. S7**). Previous studies have reported strong associations between lipid-bound α-syn species and PD pathology, raising the possibility that some assemblies visualized by Syn-IMAGR may arise from clustering of lipid-associated α-syn monomers rather than protein-only aggregates. Although we cannot fully exclude this possibility, direct evaluation of lipid-associated α-syn within the current Syn-IMAGR workflow remains technically challenging due to the absence of compatible lipid staining alongside α-syn imaging. Annexin V, a phosphatidylserine-binding probe commonly used for membrane and extracellular vesicle labeling, was used to assess whether α-syn aggregates were associated with lipid-containing structures (*34, 35*). However, in dual-color imaging experiments using Annexin V and anti-α-syn antibodies on brain lysates, we observed apparent interactions between Annexin V staining and α-syn antibody labeling, complicating interpretation of lipid-associated species (data not shown).

This behavior was further clarified by fixation experiments. Once α-syn species were chemically cross-linked, they resisted dissociation during the multiple wash steps required in the Syn-IMAGR workflow, allowing a greater fraction of aggregates—especially PD-associated aggregates—to remain detectable. This observation is consistent with previous reports showing that *in vitro*–synthesized α-syn oligomers become dissociation-resistant following fixation (*36*). Conversely, at 1,000-fold dilution, cross-linking prevented physiological multimers from naturally disassembling into smaller species, thereby reducing the apparent number of total aggregates. Together, these findings show that fixation differentially stabilizes α-syn species and prevents them from dissociating, revealing that physiological multimers are highly labile, whereas PD-associated aggregates remain stable and detectable even without fixation.

In CSF, disease-associated differences were largely masked by extreme dilution. Aggregate counts appeared similar between PD and controls, but PD CSF contained markedly lower total α-syn, resulting in reduced aggregate-to-total ratios. Importantly, CSF is several orders of magnitude more dilute than brain lysate; under these conditions, physiological multimers and small assemblies are expected to dissociate below the detection threshold, thereby masking differences in aggregate stability that are readily observed in brain tissue. Thus, the apparent similarity in CSF aggregate counts likely reflects dilution-driven loss of detectable assemblies, rather than a true absence of group differences.

The equilibrium model revealed by brain and CSF measurements aligns with calibration behavior. The antibody B/B pair displayed a characteristic concentration-dependent peak at ∼0.16 pg/mL, followed by reduced signal at higher dilutions—reflecting partial disassembly of large aggregates into smaller species. This phenomenon, analogous to dilution-induced destabilization in seed amplification assays, underscores that α-syn aggregate stability is conditional. Similarly, preanalytical variables such as freeze–thaw cycles may further perturb aggregate structure and equilibrium distributions, potentially contributing to variability across studies. Quantifying spot intensity and abundance across serial dilutions therefore allows Syn-IMAGR to infer population shifts across the free-energy landscape connecting monomers, oligomers, and higher-order assemblies.

Several technical refinements could further enhance the mechanistic resolution of Syn-IMAGR. First, the detection antibody B, is labeled using NHS-dye chemistry, resulting in conjugation of multiple dyes to each antibody. While a higher degree of labeling improves the signal-to-noise ratio, it complicates the interpretation of spot intensity in terms of antibody binding stoichiometry. Site-specific labeling strategies would enable more accurate quantification of antibody occupancy per aggregate and, by extension, more precise estimates of the number of α-syn molecules per assembly. Second, measurements of α-syn aggregates in biofluids differ from measurements obtained using calibration standards due to the heterogeneous mixture of α-syn species present in biofluids. Since antibody B captures all α-syn forms, competition for surface binding may reduce apparent aggregate abundance and lead to underestimation. Third, biological samples with limited extracellular dilution, such as brain lysates or skin biopsies, may be more suitable for detecting PD-associated aggregates, as physiological multimers are less likely to dissociate below detection thresholds. Fourth, because dilution perturbs the equilibrium of physiological α-syn species, optimizing dilution conditions for each sample type will be essential for accurate quantification. While mild cross-linking can be used experimentally to stabilize assemblies and minimize dissociation during measurement, preserving native equilibrium behavior is advantageous for distinguishing transient from stable species. Finally, we did not systematically investigate the effect of buffer composition on aggregate stability. Buffers with lower ionic strength may better preserve aggregate integrity during dilution (*37, 38*). Systematic exploration of these parameters, together with the development of higher-affinity and more selective binding reagents, would further improve Syn-IMAGR’s ability to resolve distinct α-syn species across the equilibrium landscape.

## Conclusion

In conclusion, Syn-IMAGR enables sub-femtomolar, single-molecule resolution of α-syn species across biological matrices. By screening multiple binders, we identified antibody B/B as the most reliable pair for detecting aggregated α-syn and showed that α-syn assemblies exist in a dynamic, concentration-dependent equilibrium rather than as static fibrils. In PD brain lysates, this equilibrium is shifted toward more abundant and structurally stable aggregates, reflected by increased aggregate-to-total α-syn ratios. In CSF, extreme dilution drives physiological multimers below detection thresholds, masking disease-associated differences that are readily observed in brain tissue. Together, these results recast PD aggregation as a metastable reorganization of α-syn homeostasis and establish Syn-IMAGR as a quantitative platform for interrogating protein aggregation equilibria in health and disease.

## Figure

**Figure S8.**
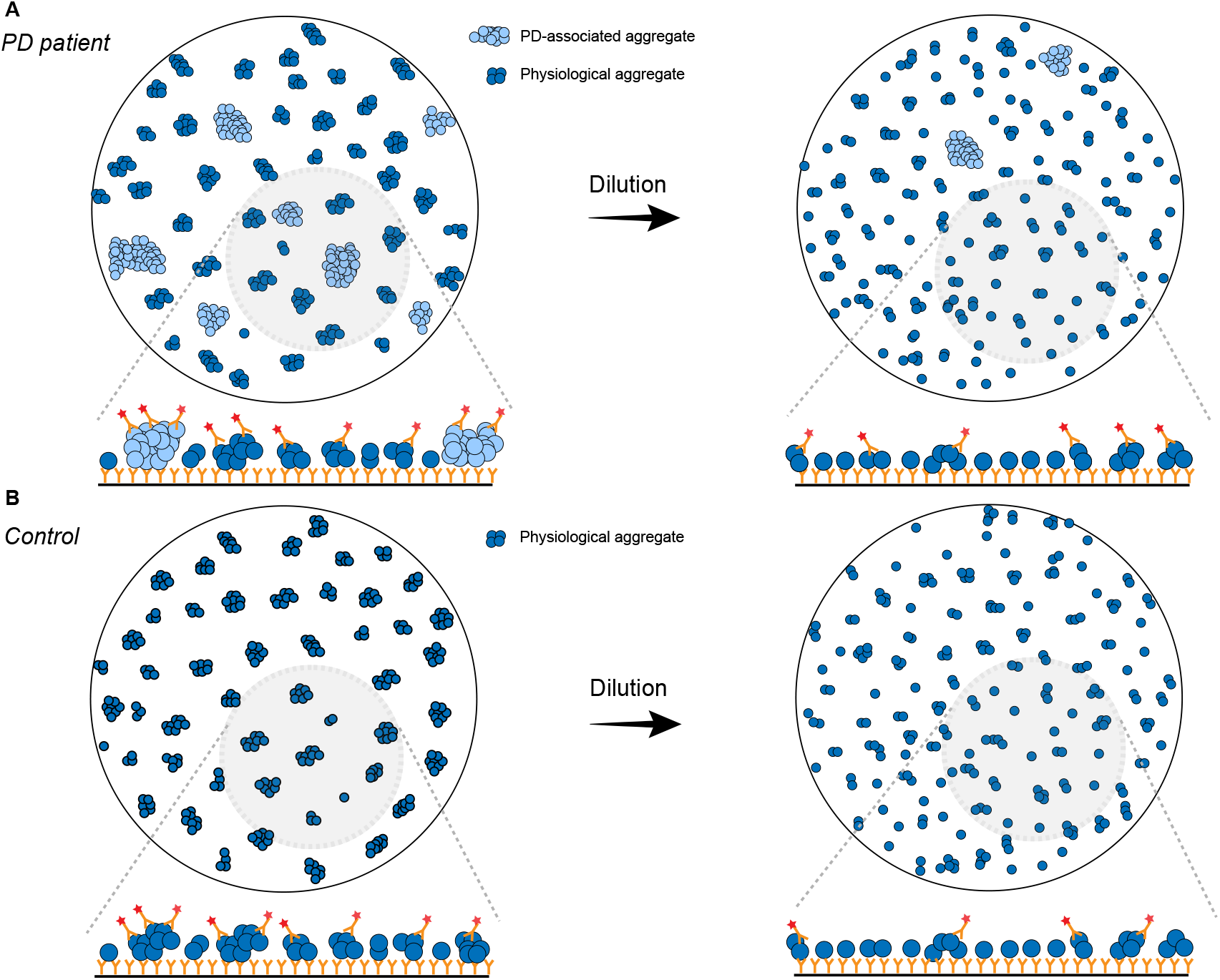
Dilution alters detectable α-synuclein assemblies differently in PD and control brain lysates. (A) *PD brain lysate*. Before dilution, PD samples contain abundant, structurally stable α-syn aggregates. After dilution, PD-associated aggregates largely remain detectable, reflecting their resistance to dissociation and disruption. (B) *Neurological control lysate*. Control samples contain predominantly physiological multimers held together by reversible, concentration-dependent interactions. Upon dilution, these multimers dissociate into smaller species that fall below detection, resulting in a marked loss of detectable aggregates at the surface.

## References

1. C. Marras et al., Prevalence of Parkinson’s disease across North America. Npj Parkinsons Dis 4, (2018).

2. R. T. Scheife, G. T. Schumock, A. Burstein, M. D. Gottwald, M. S. Luer, Impact of Parkinson’s disease and its pharmacologic treatment on quality of life and economic outcomes. Am J Health-Syst Ph 57, 953–962 (2000).

3. A. W. Willis et al., Incidence of Parkinson disease in North America. Npj Parkinsons Dis 8, (2022).

4. T. Pringsheim, N. Jette, A. Frolkis, T. D. L. Steeves, The Prevalence of Parkinson’s Disease: A Systematic Review and Meta-analysis. Movement Disord 29, 1583–1590 (2014).

5. M. Politis, G. Pagano, F. Niccolini, Imaging in Parkinson’s Disease. Int Rev Neurobiol 132, 233–274 (2017).

6. G. Pagano, F. Niccolini, M. Politis, Imaging in Parkinson’s disease. Clin Med 16, 371–375 (2016).

7. P. Mahlknecht et al., Optimizing odor identification testing as quick and accurate diagnostic tool for Parkinson’s disease. Movement Disord 31, 1408–1413 (2016).

8. A. P. Zarzur, A. D. Duprat, B. O. Cataldo, D. Ciampi, E. Fonoff, Laryngeal Electromyography as a Diagnostic Tool for Parkinson’s Disease. Laryngoscope 124, 725–729 (2014).

9. H. Golan, O. Volkov, E. Shalom, Nuclear imaging in Parkinson’s disease: The past, the present, and the future. J Neurol Sci 436, (2022).

10. R. Raghunathan, K. Turajane, L. C. Wong, Biomarkers in Neurodegenerative Diseases: Proteomics Spotlight on ALS and Parkinson’s Disease. Int J Mol Sci 23, (2022).

11. R. Chen, X. Gu, X. Y. Wang, α-Synuclein in Parkinson’s disease and advances in detection. Clin Chim Acta 529, 76–86 (2022).

12. A. Atik, T. Stewart, J. Zhang, Alpha-Synuclein as a Biomarker for Parkinson’s Disease. Brain Pathol 26, 410–418 (2016).

13. M. Fayyad et al., Parkinson’s disease biomarkers based on α-synuclein. J Neurochem 150, 626–636 (2019).

14. P. G. Foulds et al., A longitudinal study on α-synuclein in blood plasma as a biomarker for Parkinson’s disease. Sci Rep-Uk 3, (2013).

15. B. Dehay et al., Targeting α-synuclein for treatment of Parkinson’s disease: mechanistic and therapeutic considerations. Lancet Neurol 14, 855–866 (2015).

16. H. R. Morris, M. G. Spillantini, C. M. Sue, C. H. Williams-Gray, Parkinson’s Disease 2 The pathogenesis of Parkinson’s disease. Lancet 403, 293–304 (2024).

17. C. Soto, α-Synuclein seed amplification technology for Parkinson’s disease and related synucleinopathies. Trends Biotechnol 42, 829–841 (2024).

18. D. Coughlin et al., α-Synuclein Seed Amplification Assay Amplification Parameters and Progression in Parkinson’s disease (PD). Movement Disord 39, S46–S46 (2024).

19. H. H. Tsao, C. G. Huang, Y. R. Wu, Detection and assessment of alpha-synuclein in Parkinson disease. Neurochem Int 158, (2022).

20. L. B. Lassen et al., ELISA method to detect α-synuclein oligomers in cell and animal models. Plos One 13, (2018).

21. T. Gilboa et al., Measurement of α-synuclein as protein cargo in plasma extracellular vesicles (vol 121, e2408949121, 2024). P Natl Acad Sci USA 122, (2025).

22. S. J. Zhang, C. Wu, D. R. Walt, A Multiplexed Digital Platform Enables Detection of Attomolar Protein Levels with Minimal Cross-Reactivity. Acs Nano 18, 29891–29901 (2024).

23. S. T. Kumar et al., How specific are the conformation-specific α-synuclein antibodies? Characterization and validation of 16 α-synuclein conformation-specific antibodies using well-characterized preparations of α-synuclein monomers, fibrils and oligomers with distinct structures and morphology. Neurobiol Dis 146, (2020).

24. B. Fauvet et al., α-Synuclein in Central Nervous System and from Erythrocytes, Mammalian Cells, and Exists Predominantly as Disordered Monomer. J Biol Chem 287, 15345–15364 (2012).

25. K. Schwab et al., Solubility of α-synuclein species in the L62 mouse model of synucleinopathy. Sci Rep-Uk 14, (2024).

26. F. Llorens et al., Validation of α-Synuclein as a CSF Biomarker for Sporadic Creutzfeldt-Jakob Disease. Mol Neurobiol 55, 2249–2257 (2018).

27. A. Mobed, S. Razavi, A. Ahmadalipour, S. K. Shakouri, G. Koohkan, Biosensors in Parkinson’s disease. Clin Chim Acta 518, 51–58 (2021).

28. R. S. Saleeb et al., Two-color coincidence single-molecule pulldown for the specific detection of disease-associated protein aggregates. Sci Adv 9, (2023).

29. C. P. Mao et al., Protein detection in blood with single-molecule imaging. Sci Adv 7, (2021).

30. Y. P. Zhang et al., Imaging Protein Aggregates in Parkinson’s Disease Serum Using Aptamer-Assisted Single-Molecule Pull-Down(Vol,95,Pg,15254-15263,2023). Anal Chem 95, 17954–17955 (2023).

31. N. M. Jensen et al., MJF-14 proximity ligation assay detects early non-inclusion alpha-synuclein pathology with enhanced specificity and sensitivity. Npj Parkinsons Dis 10, (2024).

32. J. Zamel et al., Structural and dynamic insights into α-synuclein dimer conformations. Structure 31, (2023).

33. T. Bartels, J. G. Choi, D. J. Selkoe, alpha-Synuclein occurs physiologically as a helically folded tetramer that resists aggregation. Nature 477, 107–110 (2011).

34. N. Arraud, C. Gounou, D. Turpin, A. R. Brisson, Fluorescence triggering: A general strategy for enumerating and phenotyping extracellular vesicles by flow cytometry. Cytometry A 89, 184–195 (2016).

35. B. Gyorgy et al., Detection and isolation of cell-derived microparticles are compromised by protein complexes resulting from shared biophysical parameters. Blood 117, e39–48 (2011).

36. H. Ruesink et al., Stabilization of α-synuclein oligomers using formaldehyde. Plos One 14, (2019).

37. M. Ziaunys, A. Sakalauskas, K. Mikalauskaite, V. Smirnovas, Polymorphism of Alpha-Synuclein Amyloid Fibrils Depends on Ionic Strength and Protein Concentration. Int J Mol Sci 22, (2021).

38. M. A. Saraiva, M. H. Florêncio, Early α-synuclein aggregation is overall delayed and it can occur by a stepwise mechanism. Biochem Bioph Res Co 635, 30–36 (2022).

